# Heritability of individualized cortical network topography

**DOI:** 10.1101/2020.07.30.229427

**Authors:** Kevin M. Anderson, Tian Ge, Ru Kong, Lauren M. Patrick, R. Nathan Spreng, Mert R. Sabuncu, B.T. Thomas Yeo, Avram J. Holmes

## Abstract

Human cortex is patterned by a complex and interdigitated web of large-scale functional networks. Recent methodological breakthroughs reveal variation in the size, shape, and spatial topography of cortical networks across individuals. While spatial network organization emerges across development, is stable over time, and predictive of behavior, it is not yet clear to what extent genetic factors underlie inter-individual differences in network topography. Here, leveraging a novel non-linear multi-dimensional estimation of heritability, we provide evidence that individual variability in the size and topographic organization of cortical networks are under genetic control. Using twin and family data from the Human Connectome Project (n=1,023), we find increased variability and reduced heritability in the size of heteromodal association networks (*h*^*2*^: M=0.33, SD=0.071), relative to unimodal sensory/motor cortex (*h*^*2*^: M=0.44, SD=0.051). We then demonstrate that the spatial layout of cortical networks is influenced by genetics, using our multi-dimensional estimation of heritability (*h*^*2*^*-*multi; M=0.14, SD=0.015). However, topographic heritability did not differ between heteromodal and unimodal networks. Genetic factors had a regionally variable influence on brain organization, such that the heritability of network topography was greatest in prefrontal, precuneus, and posterior parietal cortex. Taken together, these data are consistent with relaxed genetic control of association cortices relative to primary sensory/motor regions, and have implications for understanding population-level variability in brain functioning, guiding both individualized prediction and the interpretation of analyses that integrate genetics and neuroimaging.

**Significance:** The widespread use of population-average cortical parcellations has provided important insights into broad properties of human brain organization. However, the size, location, and spatial arrangement of regions comprising functional brain networks can vary substantially across individuals. Here, we demonstrate considerable heritability in both the size and spatial organization of individual-specific network topography across cortex. Genetic factors had a regionally variable influence on brain organization, such that heritability in network size, but not topography, was greater in unimodal relative to heteromodal cortices. These data suggest individual-specific network parcellations may provide an avenue to understand the genetic basis of variation in human cognition and behavior.

## Introduction

The cerebral cortex is organized into a tightly interdigitated set of large-scale functional networks. Seminal tract-tracing work in non-human primates first revealed the structural properties underlying the distributed and parallel organization of cortical networks^1^. Subsequent resting-state functional connectivity magnetic resonance imaging (fcMRI) analyses leveraged correlation patterns of intrinsic fMRI signal fluctuations in humans^2^ to establish a canonical network architecture that is broadly shared across the population^3–8^. Yet, many individual-specific properties of brain network organization are lost when central tendencies are examined across large groups. The use of population-average network topographies has accelerated psychological and neuroscientific discovery, however there is growing recognition that the human brain is characterized by striking functional variability across individuals^9–15^. As individualized approaches become increasingly popular for the study of human behavior and psychopathology^13,16–18^, there is growing need to quantify the heritable bases of population-level variability in functional network size and topography. Despite the fact that individual differences result from the convergence of both genetic and environmental influences, the extent to which the size and spatial patterning of cortical networks may reflect heritable features of brain function has not yet been systematically investigated.

Population-based neuroimaging studies have revealed core principles that govern the evolution^19^, development^20^, and organization^7,8^ of large-scale brain networks. In particular, fcMRI has been widely utilized to generate group-average network templates through the joint analyses of data across vast numbers of individuals. The topography of these population-based network solutions are closely coupled to cognitive function, and a strong correspondence has been observed linking the spatial structure of intrinsic (fcMRI) and extrinsic (task-evoked) networks of the human brain^21–23^. Consistent with these observations, various connectivity patterns track behavioral variability in the general population^24–26^ and symptom expression in patients with psychiatric illness^27^. Suggesting genetic factors may influence the functioning of large-scale brain networks, patterns of intrinsic connectivity within population-average defined network templates are heritable^28–30^ and act as a trait-like fingerprint that can accurately identify specific people from a larger group^31,32^. These data have provided the empirical scaffolding necessary to examine how genetic, molecular, and cellular mechanisms shape human brain function^33–35^. Critically however, the use of population-based network templates can obscure individual-specific features of brain organization^9^, and there is growing evidence for substantial inter-individual variability in the size, location, and topographic arrangement of regions comprising spatially distributed functional networks across the cortical sheet.

The presence of individual differences in connectome organization presents a challenge for neuroscientists studying the functional architecture of the human brain. The identification of genetic and developmental cascades that underpin population-level variability in brain function is partly dependent on whether a group network atlas aligns to the particular functional topography of an individual. As one example, reports of population-level variability in connectivity strengths across participant groups may in fact emerge from the misalignment of underlying functional networks, obscuring the distinction between individual differences in network connectivity and topography^36,37^. Moreover, personalized network parcellations may be preferable for predictive modeling, graph theoretic, and imaging genetic approaches where the definition of an areal “unit” of cortex can influence downstream interpretations^38^. While the size and shape of individualized networks are stable across time^39^, predictive of behavior^13,40^ and refined over the course of development^41^, the molecular and genetic bases of this variability in network size, location, and spatial arrangement remain to be established. Recent studies have shown that individual differences in functional connectivity are heterogeneous across the cortex, with greater variability in association cortex relative to unimodal regions^13,15,39,42^. This distribution may have practical implications for the heritability of network topographies in association cortices, pointing to potential relationships linking the spatial distribution of inter-individual variation in functional connectivity, brain evolution, and development. Quantifying the heritability of individual-specific network topographies across the cortical sheet could provide new insights into the biological underpinnings of individual differences in human brain functions.

Although prior twin studies establish the heritability of functional connectivity strength within population-average network templates^28,29,43,44^, the role of genetics in sculpting the spatial topography of the functional connectome has yet to be quantified. To directly address this open question, we couple a multi-session hierarchical Bayesian model (MS-HBM) for estimating individual specific cortical networks^13^ with a novel non-linear multi-dimensional estimation of heritability. This approach allows us to establish the extent to which genetic and environmental factors influence individual differences in network size and topography across the cortical sheet. In doing so we provide evidence that inter-individual variability in both the spatial extent and topographic organization of cortical networks are under genetic control.

## Results

### Inter-individual variability in network sizes is nonuniform across heteromodal and unimodal cortices

We first characterized inter-individual variability in the size of functional networks across the cortical sheet. Individual-specific network topographies for each HCP participant were obtained from the method of Kong and colleagues^13^, derived using a multi-session hierarchical Bayesian model (MS-HBM). For every participant, each vertex on the cortical surface was assigned to one of 17 canonical functional networks^8^, based on both intra-individual and inter-individual patterns of cortical resting-state correlation (Figure 1a). Networks were broadly divided into those encompassing unimodal sensory and motor regions (i.e. Visual A/B/C, Somato/motor A/B, and Auditory), and those linked to heteromodal association cortex (i.e. Default A/B/C, Control A/B/C, Ventral Attention A/B, Dorsal Attention A/B, and Language). HCP cortical parcellations were identical to those of Kong and colleagues^13^, who first detailed the MS-HBM approach and demonstrated that individualized network topographies are predictive of behavior. Parcellations were derived from surface-based rsfMRI data aligned to a surface-mesh group template (fs_LR32k). For each participant, we masked out the midline and generated a 59,412 vertex array of network labels, where each vertex is assigned to one of 17 networks.

**Figure 1.**
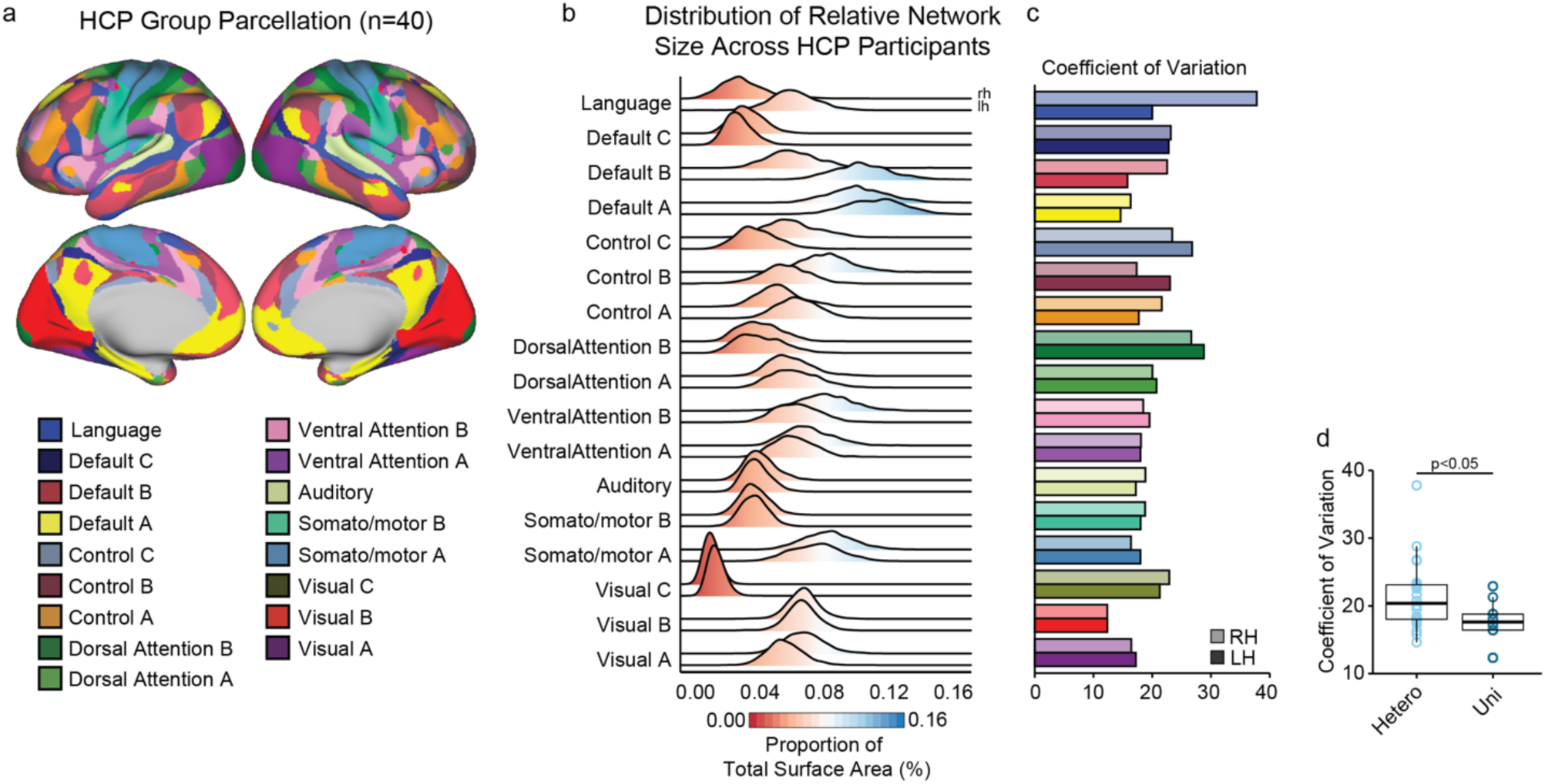
Individualized network size is more variable in heteromodal relative to unimodal cortex. (a) Individualized parcellations are composed of 17 canonical functional networks present in all HCP individuals, as defined by Kong and colleagues^13^. (b) The relative size of individualized networks was calculated for each participant, expressed as a fraction of total cortical surface area. The ridge plot shows distributions of network size across all individuals, separated by hemisphere (top ridge=right hemisphere, bottom ridge=left hemisphere). (c) Variability of individualized network size across all participants, measured with coefficient of variation, which corrects for differences in the total average size of each network. (d) Network sizes are significantly more variable within heteromodal (M=21.5, SD=5.19) relative to unimodal cortices (M=17.5, SD=3.06; F(1,32)=6.03, p=0.019). Hetero, heteromodal cortex; Uni, unimodal cortex; RH, right hemisphere; LH, left hemisphere.

The spatial extent of each network within an individual was estimated as the summed surface area of all network labeled vertices, derived using each individual’s Freesurfer-estimated vertex surface area. Differences in total cortical size across participants were adjusted by dividing summed network area by total surface area (separately for each hemisphere), resulting in a measure of relative network size across the HCP sample. Networks displayed non-uniform patterns of variability across individuals, as displayed in Figure 1b. Between-participants variability in network size was quantified using coefficient of variation (See *Methods*), which corrects for baseline differences in average network surface area. Overall, areal size was significantly more variable among heteromodal networks relative to unimodal (F(1,32)=6.03, p=0.019), an effect that remained if we used standard deviation as a measure of variability rather than coefficient of variation (F(1,32)=21.57, p=5.57e-05). These data are in line with prior reports indicating that inter-individual variability in the strength of functional connectivity is greatest in heteromodal cortex^42^, corresponding to territories with highest evolutionary cortical expansion and density of long-range functional connections^45,46^.

### Reduced heritability of network size in heteromodal relative to unimodal cortex

Inter-individual variability in network connectivity strengths are, in part, attributable to genetic variation across the population^29^. However, the majority of the literature on the genetic bases of network architecture relies on population-level motifs derived by averaging data across large groups of spatially normalized individuals^28,30,44^. To advance our understanding of the biological bases of network organization, it is important to move from group-level parcellations to a level of granularity that is only accessible when studying network organization within the individual. Given that both the size and shape of individualized functional networks are tied to behavior^13,17,39,41^, it is critical to determine heritable sources of variation that govern the amount of cortex occupied by a given functional network.

Analyses revealed that the sizes of individualized networks were significantly heritable across all canonical large-scale functional networks (Figure 2a). Heritabilities (h^2^) were calculated using individualized network size (adjusted for total surface area) and ranged between 0.22-0.57 (M=0.37, SD=0.08). Heritability of normalized surface area for each network was estimated using SOLAR^47^, and covaried for age, age^2^, age ^⊤^ sex, age^2^ ^⊤^ sex, ethnicity, height, BMI, and Freesurfer-derived intracranial volume. Suggesting broad consistency in the influence of genetic factors on the size of cortical networks across hemispheres, a significant positive correlation between left- and right-hemisphere heritability estimates was evident across the 17 networks (Figure 2b; Pearson’s r(15)=0.74, p=7.4e-4; Spearman’s *r*_*s*_=0.71, p=0.002). Notably, heritability was significantly greater within unimodal networks (h^2^: M=0.44, SD=0.05) than networks within heteromodal (h^2^: M=0.33, SD=0.07) association cortices (Figure 2c; F(1,32)=6.03, p=5.52e-05). These data demonstrate the substantial influence of genetic factors on the spatial extent of cortical networks across individuals. The results are consistent with the hypothesis that late developing aspects of heteromodal association cortex are under relaxed genetic control relative to unimodal cortex^48^. It is important to emphasize, however, that heritability refers to genetic variance accounting for inter-individual differences in a given environmental context, not the degree to which an overall trait is evolutionary constrained or genetically encoded.

**Figure 2.**
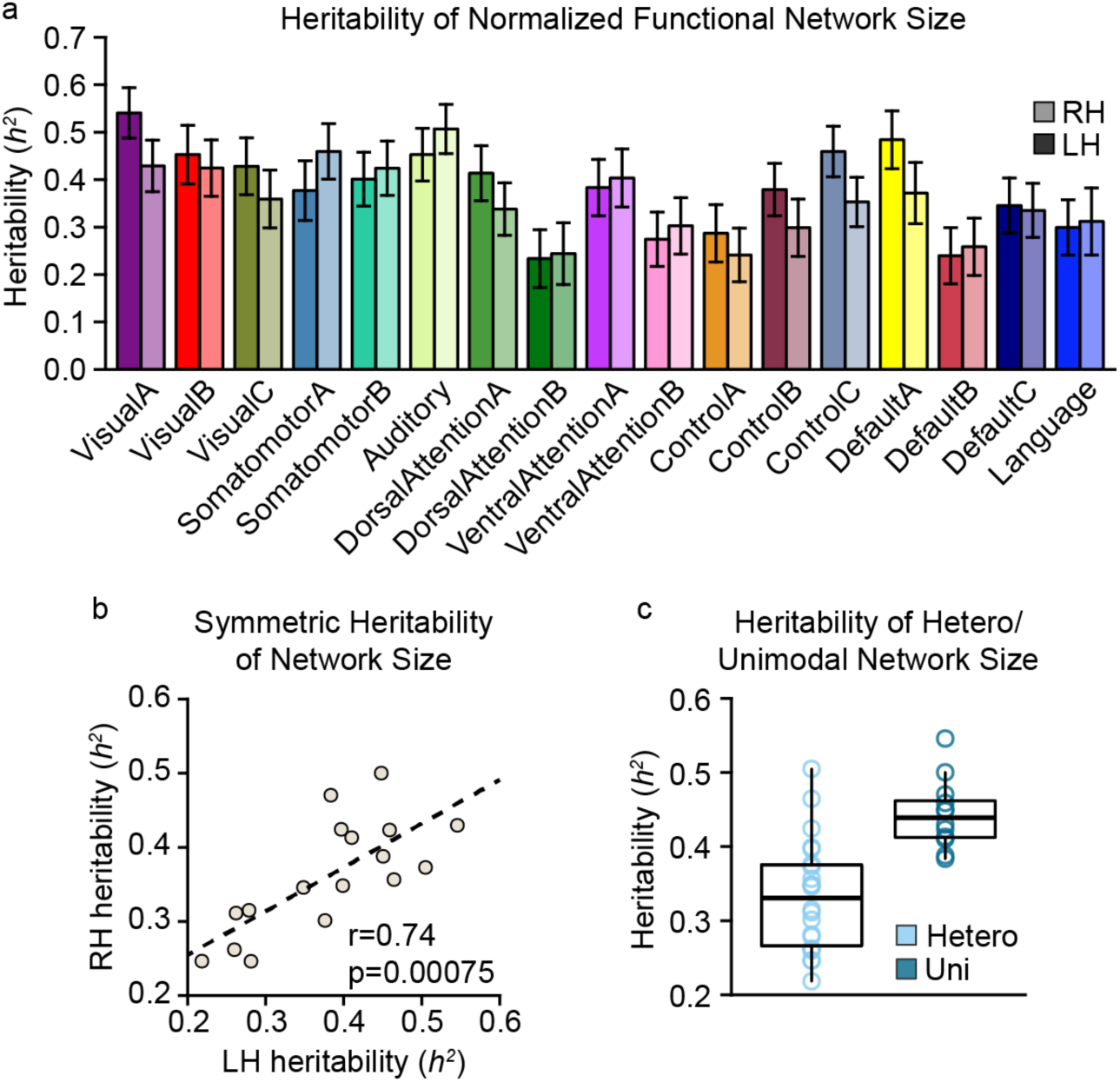
Heritability of individualized network size is greater in unimodal relative to heteromodal networks. (a) Heritability of individual network size (normalized for total surface area) was estimated across 17 canonical functional networks using SOLAR^47^, separately for each hemisphere. Error bars reflect 95% C.I. (b) The amount of variance explained by genetics (*h*^2^) for each network was consistent across hemispheres, as revealed by a correlation of left- and right hemisphere *h*^*2*^ values (r=0.74, p=7.5e-4). Each dot in the correlation plot is a functional network. (c) Heritability of normalized individual network size was higher among unimodal/sensory networks relative to heteromodal association networks (p=5.52e-05). Each dot represents one of 17 cortical networks, split by hemisphere (n=34). See Fig. 1 legend for explanation of abbreviations.

### Heritability of individualized network topography across cortex

The connectivity strength and organizational properties of functional networks vary across individuals^31,42^. Individualized parcellation approaches have established similar patterns of inter-individual variation in terms of cortical network topography, operationalized here as the spatial configuration of a given network on the cortical sheet^11,13^. A number of factors may play a role in differentiating functional topography across individuals, including mechanical tension of neuronal projections^49^, cellular and molecular properties of cortex^50^, variations in early cortical arealization by embryonic molecular patterning centers^51^, and the fundamental role sensory input plays in shaping functional organization across the cortical sheet^52^. However, the extent that variability in the spatial organization of cortical networks may be genetically driven within the general population remains unknown.

Here, we establish that genetic factors influence individualized network topographies using a novel multi-dimensional estimator of heritability. In traditional heritability analyses, the variability of a continuous (e.g. height) or categorical (e.g. diagnosis) phenotype is decomposed into the relative effects of additive genetics (A), shared environment (C), and unique environment (E; ACE model^53^). Network topography, however, is inherently multi-dimensional, since any given cortical vertex is categorically assigned to one of a set of functional networks. To account for this property of network organization we developed a novel approach to estimate heritability from a linear or nonlinear phenotypic similarity matrix defined across individuals. Inter-individual covariance of network shape was measured using Dice coefficient, which quantifies the amount of spatial overlap for any given network and participant pair (See Figure 3d for example). That is, higher Dice coefficients correspond to more similar network configurations. The observed Dice coefficients were variable across individuals, as well as non-uniformly distributed across networks (Figure 3a). The unimodal networks were overall more similar across individuals (Dice: M=0.77, SD=0.05) relative to heteromodal association networks (Dice: M=0.55, SD=0.06; F(1,32)=112.4, p=5.35e-12). The increased topographic variability of association networks is consistent with prior reports of greater inter-individual variation in accompanying patterns of long-range connectivity^42^.

**Figure 3.**
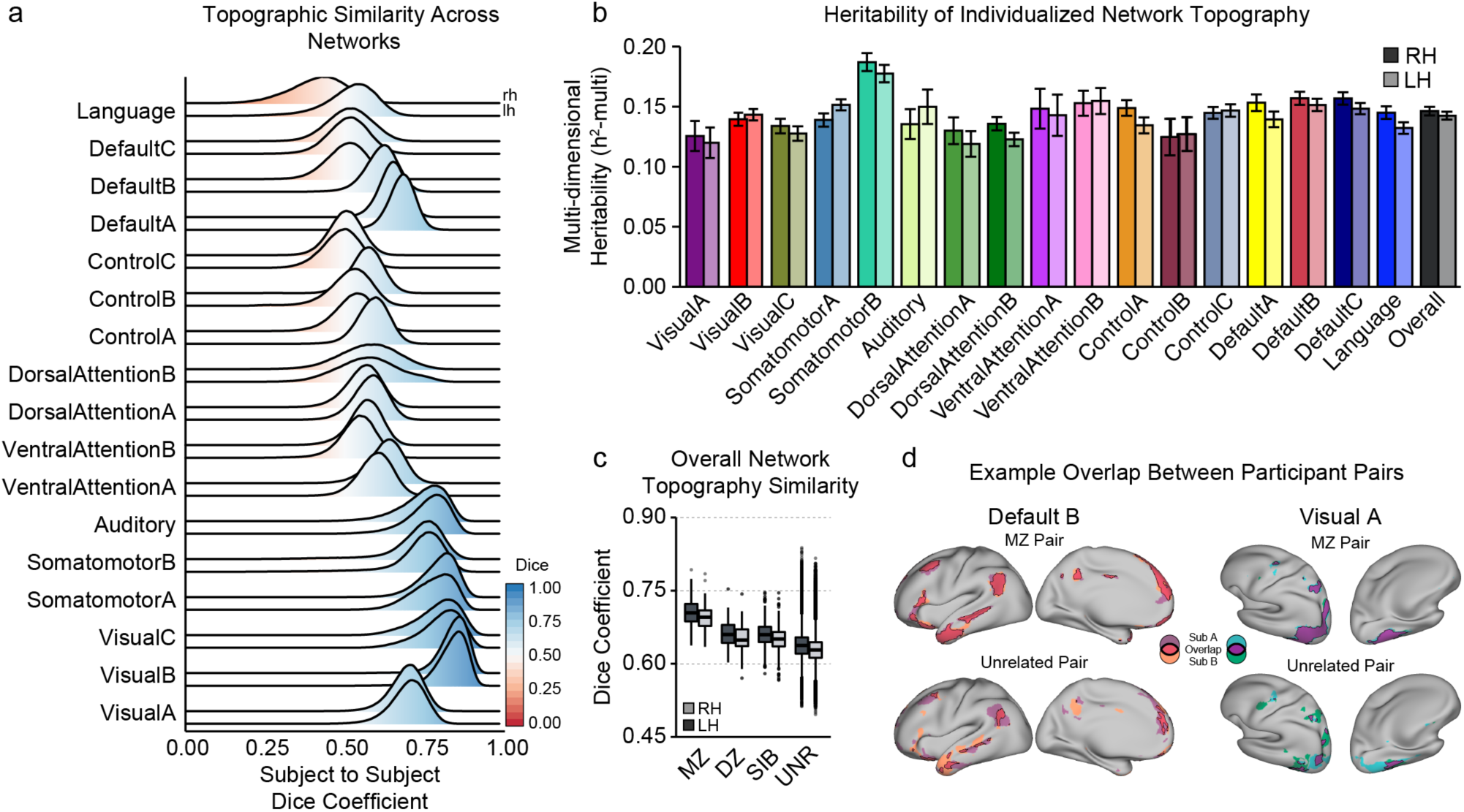
Individualized network topography is heritable across all networks. (a) The ridge plot displays distributions of inter-individual Dice coefficients across each network. Higher Dice coefficients reflect higher spatial overlap of a network for a given pair of individuals. Topography of unimodal networks were overall more similar across individuals, relative to heteromodal cortex. (b) Significant heritability was observed across all examined 17 cortical networks (q<0.01; range=0.12-0.19, mean=0.14), which was symmetric across hemispheres (r=0.84, p=2.8e-5; *r*_*s*_=0.76, p=0.0006). (c) Boxplots show higher Dice similarity of overall network organization between MZ pairs, relative to DZ, sibling, and unrelated participant pairings. (d) Individual examples illustrate HCP participants with a high and low dice overlap for Default B (high=0.78; low=0.29) and Visual C (high=0.93; low=0.59) networks. MZ, monozygotic; DZ, dizygotic; SIB, sibling; UNR, unrelated.

Analysis of multi-dimensional heritability, denoted “h^2^-multi”, demonstrated that inter-individual differences in network topography were significantly influenced by inherited genetics (Figure 3b; h^2^-multi: min=0.12, max=0.19, M=0.14, SD=0.015), after accounting for multiple testing correction (Bonferroni False-Discovery Rate correction, q’s<0.05). Figure 3c displays the distribution of Dice coefficients reflecting inter-individual similarity of network topography defined across all 17 cortical networks (i.e. “Overall” in Figure 3b). Dice similarity was greater for monozygotic twins (LH: M=0.70, SD=0.026; RH: M=0.69, SD=0.024), relative to dizygotic twins (LH: M=0.66, SD=0.028; RH: M=0.65, SD=0.028), siblings (LH: M=0.66, SD=0.026; RH: M=0.65, SD=0.028), and unrelated individuals (LH: M=0.64, SD=0.027; RH: M=0.063, SD=0.026), corresponding to a h^2^-multi of 0.142 and 0.146 for left and right hemispheres, respectively. The degree of topographic heritability for each network was consistent across hemispheres (Spearman’s rho=0.69, p=0.0036). Contrary to estimates of individualized network size, the heritability of network topographies did not differ between unimodal and heteromodal cortices (F(1,32)=0.21, p=0.65). Overall, these data advance a novel heritability estimation technique to demonstrate that the spatial organization of functional networks is influenced by genetic factors.

We next quantified local genetic control of network architecture across the cortical sheet. Our findings detailed above show that the heritability of network topography is broadly uniform across cortex when averaging within individual networks (Figure 3). Prior work, however, indicates significant spatial heterogeneity of heritable aspects of cortical anatomy^54–56^. Here, we demonstrate that genetic influences on local network topography are also spatially variable across cortex, with the greatest heritability observed within the precuneus as well as dorsal aspects of parietal, prefrontal, and posterior parietal cortices (Figure 4a). Multi-dimensional heritability estimates were calculated in the same manner as above, whereby a Dice coefficient matrix represented the participant to participant similarity of network topography. In this analysis however, we only consider individualized network labels falling within a given region of interest (ROI; radius=10 vertices) at each point on the cortical sheet. Figure 4b illustrates example participant pairs with high and low Dice coefficients for an ROI in the precuneus. Critically, some participant pairs possess almost entirely non-overlapping network assignments within a given patch of cortex (Figure 4b), highlighting the need for individualized parcellations to study of neurobiological variability across the population. Of note, we did not observe a clear dissociation between unimodal and heteromodal cortices in terms of local network heritability. That is, heritability estimates did not differ between regions canonically associated with sensorimotor networks (M=0.149, SD=0.038) relative to association networks (M=0.146, SD=0.037; F(1,32)=2.10, p=0.16). Together, these results indicate the heritable basis of network organization is variable across cortex and support further research into the biological determinants of network topography.

**Figure 4.**
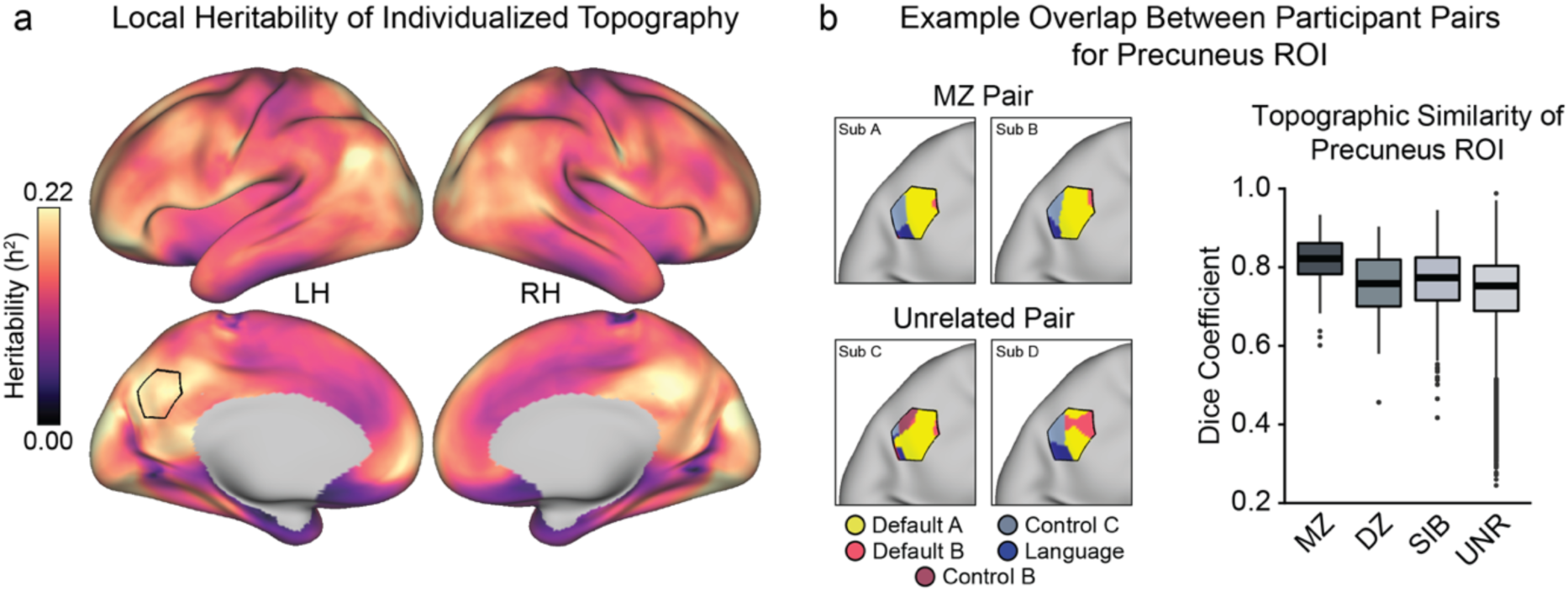
Local Heritability of Individualized Network Topography. (a) Multi-dimensional heritability of network topography estimated for every vertex, using an ROI at each point on the cortical sheet (radius=10 vertices). Individualized network labels in each ROI were evaluated to compute a participant-to-participant Dice similarity matrix, reflecting the similarity of network assignments within a given cortical area. Warmer colors indicate higher heritability of network assignments, for instance reflecting greater similarity among twins and siblings than unrelated individuals. (b) Example participant pairs with high and low Dice overlap of network labels for an ROI in the precuneus. Dice similarity was higher between MZ twins, relative to DZ, sibling, and unrelated participant pairs. MZ, monozygotic; DZ, dizygotic; SIB, sibling; UNR, unrelated; LH, left hemisphere; RH, right hemisphere.

## Discussion

The use of population-average network templates has provided foundational insights into the macroscopic functional organization of the human brain. However, individual specific features of brain network architecture are obscured when collapsing data across large groups of participants. Methodological advances make it possible to measure individualized features of functional networks *in-vivo*, promising to yield biological insight into the genetic, molecular, and cellular bases of cortical brain organization. Here, leveraging a novel form of multi-dimensional heritability analysis, we demonstrate that a substantial portion of the population-level variability in the size and spatial arrangement of cortical networks is under the influence of genetic factors. In Human Connectome Project data (n=1,023), the relative size (i.e. cortical surface area) of individualized networks showed considerable inter-individual variation, which was most pronounced in higher-order heteromodal relative to unimodal sensorimotor networks (Figure 1). We demonstrated that individualized network size was heritable for all 17 examined cortical networks, but was most pronounced within unimodal, relative to heteromodal cortices (Figure 2). Next, we established the heritability of individualized network spatial organization, or topography, for all cortical functional networks (Figure 3). Although topographic heritability was broadly consistent between cortical networks, we observed substantial spatial heterogeneity in the influence of genetic factors across the cortical sheet (Figure 4). Together, this work advances a novel analytic framework for measuring heritability of multi-dimensional traits to establish the extent that individual-specific features of functional network organization are influenced by inherited genetics.

The estimation of the heritability of multi-dimensional traits, such as the brain’s functional network architecture, is challenging given that traditional approaches are designed for continuous (e.g. height) or binary (e.g. diagnosis) phenotypes (but see^57^). In the present study, we described a novel method for estimating heritability from any matrix of participant-wise similarity metrics. Dice coefficients were used to quantify between-participants similarity of network topography, but this approach is generalizable to other commonly studied neuroscientific phenotypes, such as patterns of anatomical similarity^58^ or morphometricity^59^. We have made the associated heritability code freely available to the community (www.github.com/kevmanderson/h2_multi), along with analytic pipelines for all analyses. This work provides the basis for further elaboration of multi-dimensional heritability techniques, such as genetic correlation, that could reveal patterns of shared genetic variance with psychological phenotypes^13,41^. Individualized network parcellations also hold promise for understanding psychiatric disorders^16^, which are often heritable^60^. Identifying shared genetic substrates between individualized features of network organization and psychiatric illness will be important as individualized approaches become increasingly adopted in clinical neuroscientific research.

Here, we demonstrated that a significant proportion of the variance in network size and topography is explained by genetics, which could emerge through many possible biological pathways. For instance, the individual differences in cortical arealization that influence network organization may be determined early in neurodevelopment. That is, the cellular fate and areal identity of early cortical progenitor cells are specified in embryonic periods by spatial gradients of molecular transcription factors^61,62^, which may vary across individuals. The ability of genetic variation to shape these early developmental processes is supported by a recent Genome Wide Association Study (GWAS) documenting that common genetic polymorphisms linked to cortical surface area were enriched among regulatory elements of neural progenitor cells^63^. The topography of cortical functional networks may also be influenced by cortical neuroanatomy, such as cortical morphology and patterns of structural connectivity. In this context, biomechanical processes such as axonal tension, intracranial pressure, and the differential growth of cortical layers are thought to influence cortical folding^49,64^, which may in turn constrain the topography of functional networks. Further, the size and shape of functional network boundaries may be sculpted by thalamo-cortical connections that refine patterns of cortical arealization across development^52^. Experimentally modulating thalamic afferents can substantially impact cortical morphology and size in a pathway specific manner ^65,66^. Critically however, heritability analyses cannot disentangle the specific biological processes that influence cortical network size or topography. Rather, our data support the importance of future work utilizing statistical genetic approaches to identify the biological cascades that influence functional network topography across the cortical sheet^63^.

The emergence of analytic frameworks for capturing individualized network architectures is both a technical and theoretical advance, providing the opportunity to link cognition and behavior to population-level variability in brain organization. Landmark research has shown that individuals can be identified by patterns of whole-brain functional connectivity, conceptualized as a functional “fingerprint”^25,31^. The analogy to a fingerprint is apt. For instance, broad classes of fingerprint types exhibit a high degree of heritability, despite ridge patterns of any fingerprint being entirely unique^67^. Likewise, the brain is organized into a core functional network architecture, that nevertheless exhibits distinctive features in a trait-like manner at the level of the individual^39^. Here, we demonstrate that the distinguishing topographic features of the brain, including variability in network organization and size, are influenced by inherited genetic factors (Figure 3). Such data provide a potential “upper-bound” on the explainable variance due to additive genetic variation, and highlight the utility of future research probing the genetic mechanisms underlying inter-individual differences in functional network organization across the lifespan.

Consistent with prior work examining population average network templates, the current study provides support for the genetic bases of core features of brain function^28–30,44,68–70^. The degree of heritability for functional network size is in line with prior estimates demonstrating that ∼40-60% of the variance in within-network connectivity is explained by genetics^28,29^. Of note, we emphasize that our data reflect a snapshot of heritability estimates for a given sample in early adulthood, and that the influence of genetic factors may vary across developmental periods^71^. Perhaps counterintuitively, phenotypic heritability generally increases from childhood through adulthood, possibly reflecting genotype-environment interactions as individuals engage in behaviors that reinforce genetically influenced traits^72^. Although, there is evidence that the reverse is true in late adulthood, such that heritability decrease with age^73^. Future work should also examine whether environmental factors such as early life stress or adversity may also impact cortical network topography and connectivity^74^. Experience is also critical for the emergence of functional selectivity in some brain regions, such as the face-responsive inferotemporal cortex, which may in turn influence patterns of network connectivity and affiliation^75^. Given recent evidence of the developmentally dynamic nature of functional network organization^41^, it will be important to utilize imaging-genetic data, such as the Adolescent Brain Cognitive Development study^76^, to quantify the age-dependent influence of genetic factors on cortical network formation.

Higher-order association networks are consistently more variable than unimodal sensorimotor networks, in terms of both relative network size (Figure 1c) and topographic network similarity (Figure 3). These observations are consistent with evidence that heteromodal cortex has greater inter-individual variance in functional connectivity^42^. The increased variability of heteromodal network size coincided with lower estimates of heritability, relative to the individualized size of unimodal networks (Figure 2c). These data are in line with theories positing that late-developing aspects of cortex are more sensitive to environmental influences and extrinsic sources of network sculpting^48^. That is, higher-order networks are the most distal (or “untethered”) from both early embryonic signaling gradients and thalamus-mediated sensory inputs^48^. However, we did not observe heteromodal versus unimodal differences in heritability for measures of network topography (Figure 3-4), as we did for individualized network size.

The present study should be interpreted in light of several limitations. First, our analyses assume that participants have been brought into a common anatomical space, but we cannot rule out the role of inter-individual differences in alignment accuracy. Here, we used sophisticated surface-based alignment techniques from the HCP that rely on multi-modal areal features of cortex rather than cortical folding patterns and anatomical landmarks^77^. However, inter-individual alignment is still subject to error. We also emphasize that heritability estimates of network topography are dependent upon the accuracy and assumptions of the parcellation approach. We also note that our novel heritability estimation calculated from a linear or nonlinear phenotypic similarity matrix is equivalent to the heritability of the intrinsic multidimensional trait that generates the participant-wise phenotypic similarity (Ge et al. 2016; see Methods), and is thus different from traditional heritability analysis of a scalar phenotype (e.g., height) unless the similarity matrix is spanned by a one-dimensional vector.

In conclusion, this paper advances a novel multi-dimensional heritability technique to establish the heritability of individualized cortical functional networks, in terms of both network size and topography. We found that the size of heteromodal cortical networks was more variable and less heritable relative to unimodal networks, in line with the protracted developmental maturation of higher-order cortex that may allow for increased influence of the environment. Individualized network topography was similarly more variable among heteromodal networks, but heritability was approximately equivalent for all cortical functional networks. However, heritability analysis of local network architecture revealed a non-uniform influence of genetic factors on network organization across cortex. Together, these data establish that the size and topography of cortical functional networks are influenced by genetic factors, providing a foundation for future work disentangling the biological mechanisms that govern individual variances in brain organization.

## Methods

### Human Connectome Project

The Human Connectome Project (HCP) is a large community-based sample of twins and nuclear family members, assessed on a comprehensive set of neuroimaging and behavioral batteries. HCP analyses initially comprised 1,029 participants that were successfully processed through the individualized network parcellation approach of Kong and colleagues (Kong et al., 2019), known as the multi-session hierarchical Bayesian model (MS-HBM). HCP twin zygosity was determined by genotyped data when possible (n=410), otherwise self-report was used to identify pairs of monozygotic (MZ) and dizygotic (DZ) twins (n=76). If imaging data was not available for one twin in a pair, the usable participant was designated a “singleton” for later heritability analyses. The final sample consisted of n=1,023 individuals (n_MZ_=274; n_DZ_=160, n_not_twin_ =482, n_singleton_=107). See **Table 1** below for basic demographics across groups.

**Table 1:**
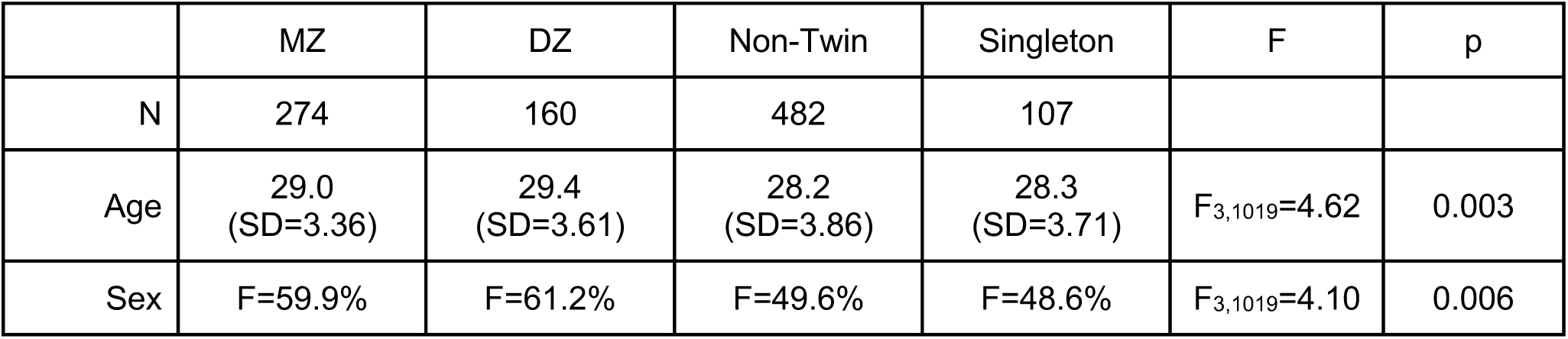
Demographics of HCP participant groups. Heritability estimates were conducted on 1,023 HCP participants, composed of 137 MZ twins (n=274), 80 DZ twins (n=160), non-twin siblings (n=482), and unrelated singletons (n=107). Although groups were nominally well-matched demographically, ANOVAs revealed significant differences of age and sex. All heritability analyses included age and sex as covariates.

### Measuring individualized network organization and size

Individualized network parcellations were derived from HCP resting-state functional magnetic resonance imaging (rs-fMRI) surface data, aligned to the *fs_LR32k* group space using the MSMAll areal-feature-based registration^78^. Methodological details of the Multi-session Hierarchical Bayesian Model (MS-HBM) approach that produced the individualized parcellations have been previously published^13^, however we present key details here. Multiband rs-fMRI data collected on Siemens 3T Skyra scanners from the HCP S1200 release were analyzed. A key feature of the MS-HBM approach is that it incorporates both intra-individual and inter-individual patterns of variability to define individualized network boundaries. HCP data is particularly suited for this method since rs-fMRI runs were collected across two separate sessions across 2 days, allowing for well-powered estimates of inter- and intra-individual variance. Individual resting state runs were 15 minutes in length and were acquired at an isotropic resolution of 2mm and TR of 0.72s^79,80^. Surface-based preprocessing of rs-fMRI data began with minimally preprocessed HCP MSMAll ICA-FIX data on a group surface template (fs_LR32k^81^). Additional preprocessing included nuisance regression, temporal censoring, and spatial smoothing.

A held-out training set of 40 HCP participants were used to derive the necessary group-level parcellation (Figure 1a) and model parameters, such as inter-individual variability, for the MS-HSM method. For each participant, each of the 59,412 bi-hemispheric vertices on the cortical sheet was assigned to one of the 17 canonical functional networks^8^. The size of each cortical network within an individual was estimated as the summed surface area of all network vertices, obtained from midthickness fs_LR32k projected Freesurfer surface area data. Network size was calculated separately for each hemisphere, then divided by total hemispheric area to quantify proportional size of a network on the cortical sheet. All cortical surface figures were created using the HCP workbench^82^.

### Heritability of Individualized Network Size

The heritability of individualized network size was estimated using SOLAR ^47^, covarying for age, sex, age^2^, age^⊤^sex, age^2⊤^sex, ethnicity, height, BMI, and Freesurfer-derived intracranial volume. Bonferroni correction of significance thresholds was used to account for 34 independent tests of heritability. Cross-hemisphere consistency (Figure 2b) was tested by correlating left- and right-hemisphere heritability estimates across all 17 cortical networks.

### Dice Similarity of Network Topography

Participant to participant similarity of individualized network topography was measured using the Dice Sørensen formula, where the coefficient for a given network reflects:

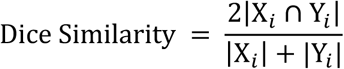

where X_*i*_ and Y_*i*_ are the network labels for network *i* for participants *X* and *Y*, ∩ represents the intersection between participant network labels, and |·| represents the total number of vertices in each set (i.e. cardinality).

### Multi-dimensional heritability analysis

Consider an *M*-dimensional trait ***Y*** = [***y***_1_, …, ***y***_*m*_] = [***y***_*im*_]_*N× M*_, and a multivariate variance component model: ***Y*** = ***G*** + ***E***, where ***G*** and ***E*** are ***N*** × ***M*** matrices representing the additive genetic effects and unique environmental factors, respectively. Assume vec(***G***)∼N(**0, ∑**_*G*_ ⊗ ***K***), vec(***E***)∼N(***0*, ∑**_*E*_ ⊗ ***I***), where vec(·) is the matrix vectorization operator that converts a matrix into a vector by stacking its columns, **⊗** is the Kronecker product of matrices, **∑**_*G*_ is the genetic covariance matrix, and **∑**_*E*_ is the environmental covariance matrix. The genetic and environmental covariance matrices can be estimated using a moment-matching method: 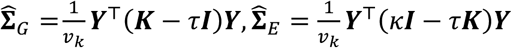 where *τ* = tr(***K***)/*N, κ* = tr(***K***^2^)/*N, v*_*K*_ = *N*(*κ* − *τ* ^2^) (Ge et al. 2016). The SNP heritability of a multidimensional trait ***Y*** is defined by 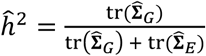 (Ge et at 2016). Note that 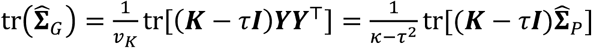, where 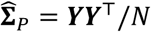 is the estimated phenotypic covariance matrix. Similarly, 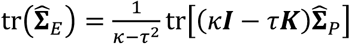. For any non-negative definite phenotypic similarity matrix 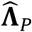 derived from a nonlinear measure, we define heritability by replacing 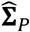 with 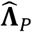:. This is known as the kernel trick in machine learning. We note that for any ***N*** × ***N*** non-negative definite matrix **∑**, there exists a ***N*** × ***P*** matrix ***V*** (often ***P*** ≪ ***N***) such that **∑** ≈ ***VV***^⊤^. Therefore, **∑** can be considered as a linear covariance matrix generated by a multidimensional trait.

To model covariates, consider a multivariate mixed effects model: ***Y*** = ***XB*** + ***G*** + ***E***, where ***X*** is an ***N*** × *q* matrix of covariates, and ***B*** is a *q* × *M* matrix of fixed effects. There exists an *N* × (*N* − *q*) matrix ***U*** satisfying ***U***^⊤^***U*** = ***I, UU***^⊤^ = ***P***_;_ = ***I*** − ***X***(***X***^⊤^***X***)^−1^*X*, and ***U***^⊤^***X*** = ***0***. Applying ***U***^⊤^to both sides of the model gives ***U***^⊤^***Y*** = ***U***^⊤^***G*** + ***U***^⊤^***E***, where vec(***U***^⊤^***G***)∼N(***0*, ∑**_*G*_⊗(***U***^⊤^***KU***)), vec(***U***^⊤^***E***)∼N(***0*, ∑**_*G*_⊗ ***I***). Therefore, we can replace ***Y*** with ***U***^⊤^***Y, K*** with ***U***^⊤^***KU***, and ***N*** with ***N*** − ***q*** in the SNP heritability estimator derived above to obtain and estimator that accounts for covariates. More specifically,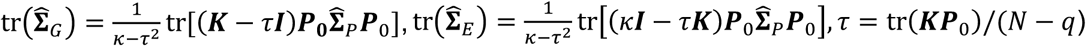, and *κ* = tr(***KP***_0_*KP*_0_)/(*N* − *q*). For nonlinear phenotypic similarity matrix 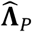:, we replace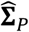: with 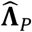:.

Significance was measured using participant-based permutations, where the kinship matrix was randomly shuffled 1,000 times. Standard errors were calculated using a block jackknife procedure with a leave one family out strategy. That is, for a given iteration of the jackknife, all participants within a nuclear family unit were excluded and heritability was re-calculated. Variability was then calculated from the resulting distribution of subsampled heritability estimates.

### Code and Data Availability

All custom code written to perform analyses are publicly available on github (https://github.com/kevmanderson/heritable_network_topography). We have also provided an open-access generalized implementation of our multi-dimensional heritability estimator (https://github.com/kevmanderson/h2_multi). Code to produce individualized MS-HBM parcellations is publicly available (https://github.com/ThomasYeoLab/CBIG). Human Connectome Project Data is available for download (https://db.humanconnectome.org).

## Acknowledgements

This work was supported by the National Institutes of Health (Grant R01MH120080 to A.J.H.; K99/R00AG054573 to T.G.; R01LM012719 and R01AG053949 to M.R.S.), the National Science Foundation (DGE-1122492 to K.M.A.; CAREER 1748377 and NeuroNex 1707312 to M.R.S.). B.T.T.Y. and R.K. are supported by the Singapore National Research Foundation (NRF) Fellowship (Class of 2017). Any opinions, findings and conclusions or recommendations expressed in this material are those of the author(s) and do not reflect the views of National Research Foundation, Singapore. Analyses were made possible by the high-performance computing facilities provided through the Yale Center for Research Computing. Data were provided [in part] by the Human Connectome Project, WU-Minn Consortium (Principal Investigators: David Van Essen and Kamil Ugurbil; 1U54MH091657) funded by the 16 NIH Institutes and Centers that support the NIH Blueprint for Neuroscience Research; and by the McDonnell Center for Systems Neuroscience at Washington University.

## Supplementary Material

**Supplementary Figure 1.**
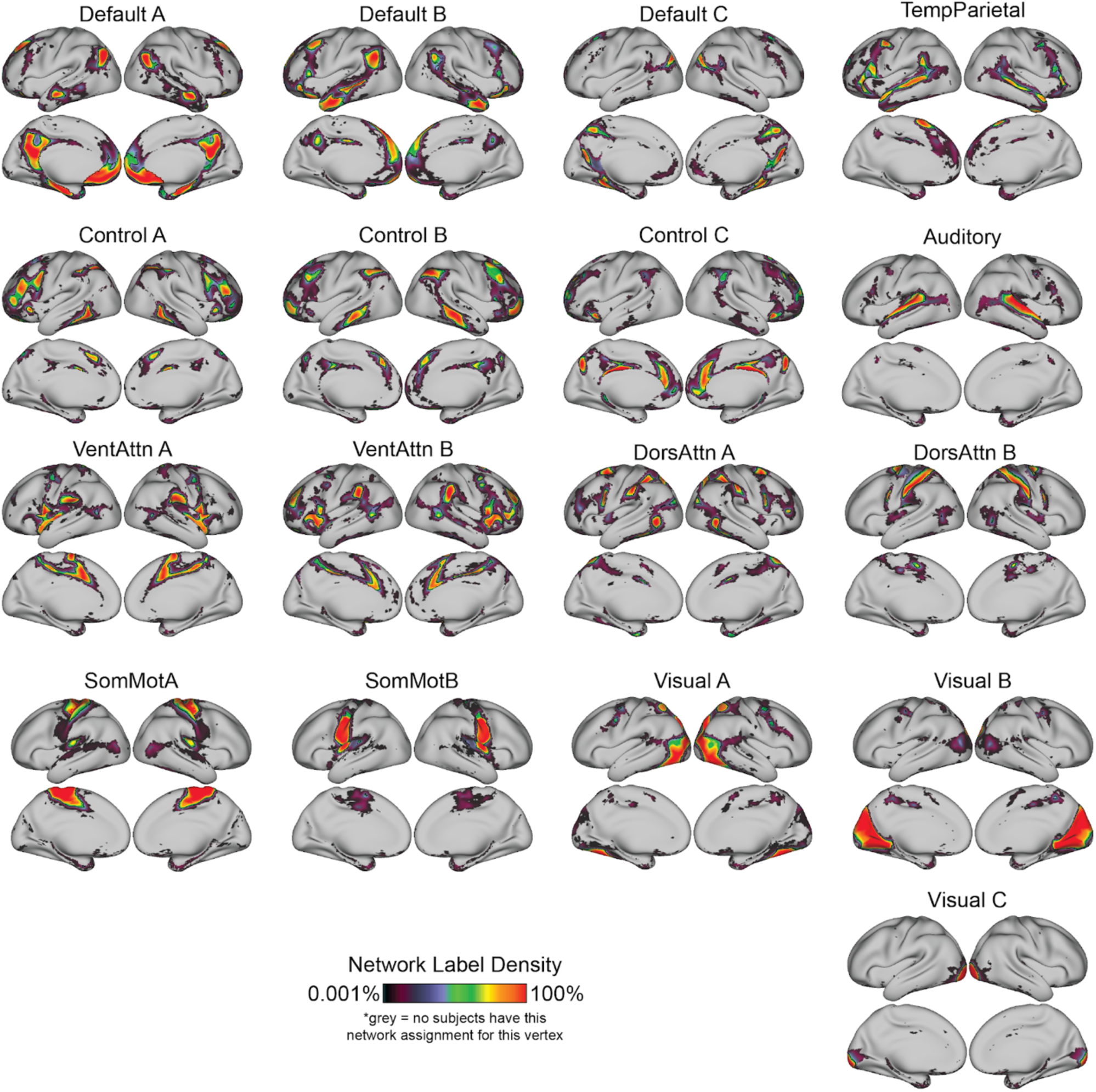
Density of individualized network topography across the cortical sheet. At each vertex, we plot the proportion of individuals that are assigned to a given network (n=1,023). Warm red indicates that a vertex is assigned to a given network in a large percentage of participants. Darker purple/black identifies cortical territories with more variable network assignment across participants. Black borders outline territories where a given network is most common (i.e. highest modal network assignment at a given vertex).

